# Unsupervised discovery of tissue architecture in multiplexed imaging

**DOI:** 10.1101/2022.03.15.484534

**Authors:** Junbum Kim, Samir Rustam, Juan Miguel Mosquera, Scott H. Randell, Renat Shaykhiev, André F. Rendeiro, Olivier Elemento

**Author notes:** Co-senior authors (Renat Shaykhiev, André F. Rendeiro, Olivier Elemento).

## Abstract

Multiplexed imaging and spatial transcriptomics enable highly resolved spatial characterization of cellular phenotypes, but still largely depend on laborious manual annotation to understand higher-order patterns of tissue organization. As a result, higher-order patterns of tissue organization are poorly understood and not systematically connected to disease pathology or clinical outcomes. To address this gap, we developed UTAG, a novel method to identify and quantify microanatomical tissue structures in multiplexed images without human intervention. Our method combines information on cellular phenotypes with the physical proximity of cells to accurately identify organ-specific microanatomical domains in healthy and diseased tissue. We apply our method to various types of images across physiological and disease states to show that it can consistently detect higher level architectures in human organs, quantify structural differences between healthy and diseased tissue, and reveal tissue organization patterns with relevance to clinical outcomes in cancer patients.

## Introduction

The recent development of technologies such as multiplexed imaging^1–5^ and spatial transcriptomics^6–10^ allow for both direct observation of cellular phenotypes and cellular interactions in native tissue microenvironment. While these technologies provide a highly resolved view of cellular heterogeneity in native tissues, they struggle to move beyond a cellcentric view of tissue, failing to uncover organizing principles of tissue architecture and tissue-specific physiology which are encoded at various scales of cellular and extracellular interactions. Understanding higher-level patterns of tissue and organ organization would be crucial to establishing a relationship between cellular phenotypes and organ-specific tissue physiology.

Visual inspection of histopathological images of biopsied or surgically removed tissue is a major component of disease diagnosis, but is a labor intensive job that requires manual annotations from specialized pathologists. Also, the process may require multiple specialists to reduce intra- and inter-observer variability. To assist and improve upon the inspection process, computational techniques have been developed for the automated detection and quantification of cells or tissue structures^11–13^, often in a supervised manner which requires manual annotations as training data. This approach is expensive and laborious, is prone to learning biases from training data, and is hard to employ with exceptionally abundant tissue features such as individual ducts in submucosal glands, or small capillaries. Unsupervised methods try to accomplish similar tasks without the need for manual input. A popular method is the inference of cell neighborhoods based on multiplexed data by assembling a graph of cellular interactions based on physical proximity^14,15^. Clustering of cells based on these interactions yields cellular neighborhoods predictive of patient survival^14–17^. However, graph clustering *per se* does not make use of cell type identities or phenotypes, and has only been applied to cancer tissue.

Recent studies applying unsupervised deep learning models to histopathological images such as hematoxylin eosin staining have shown that it is possible to extract morphological features that are for example predictive of gene expression^18^. Other studies have also employed deep learning of graphs of cellular proximity with cellular phenotypes for cell type prediction^19^, inference of cellular communication^20^, and data exploration^21^. These models are computationally expensive to train, and their results heavily depend on training data, which may preclude joint analysis of expression and morphological features across studies and data types. There is thus a need for unsupervised, broadly applicable methods of tissue structure detection across organs and imaging modalities that incorporate cellular proximity, expression, and morphological features. Here we present a novel and accurate method to perform discovery and quantification of microanatomical tissue structures in multiplexed histopathological images without human intervention or prior knowledge. Our method combines information on cellular morphology and expression with the physical proximity of cells to discover domains of tissue architecture. We demonstrate that our approach is able to discover organ-specific microanatomical domains in the human lung, across diseased physiological states, and even to cancer contexts where it uncovers tissue organization with relevance to the clinical outcome of patients.

## Results

### Unsupervised identification of tissue microanatomical domains with graphs (UTAG)

To address the problem of discovery of microanatomical structure in tissue across data types and biological systems we developed a method called <u>u</u>nsupervised discovery of <u>t</u>issue <u>a</u>rchitecture with <u>g</u>raphs (UTAG) (**Figure 1a**). Our method is generally applicable to images of cells in their native tissue context collected via highly multiplexed single-cell imaging data such as co-detection by indexing (CODEX), cyclic immunofluorescence (CyCIF), imaging mass cytometry (IMC), multiplexed ion beam imaging (MIBI), and likewise multiplexed spatial platforms. The central aspect of UTAG is the combination of two matrices that represent phenotypic and positional information about each cell present in an image (**Figure 1a,** gray areas), to generate a new feature space which encodes spatially aggregated phenotypic information. This matrix of new features can then be clustered into domains of cells that are both phenotypically and spatially related (**Figure 1a,** orange area). The matrix of phenotypic information (feature matrix) is a numeric matrix of gene or protein abundance, or morphology for each cell, while the positional information of each cell is used to generate a graph of physical proximity between cells through binarization and optional normalization (adjacency matrix).

**Figure 1:**
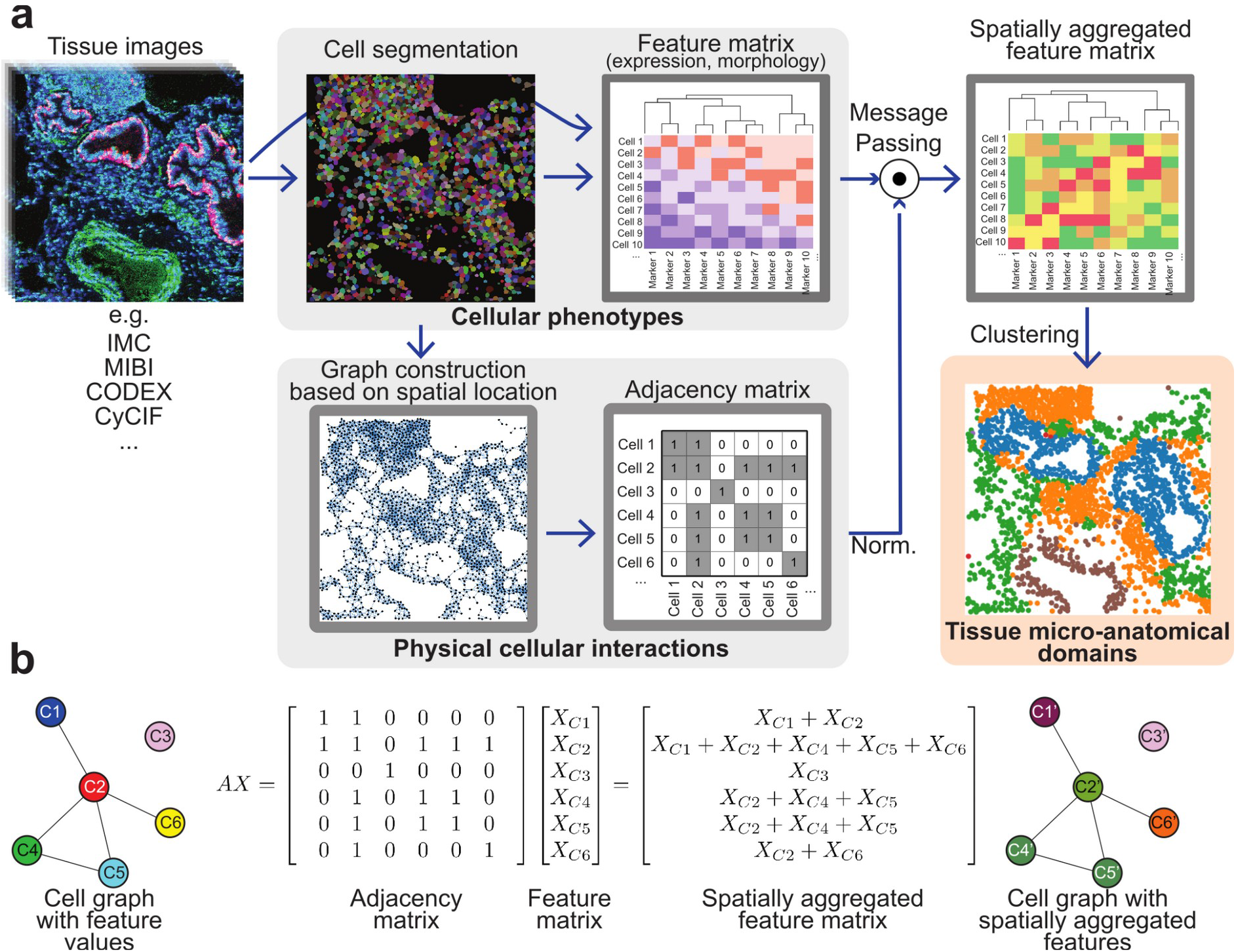
Unsupervised discovery of tissue architecture with graphs. **a**) Schematic description of the methodology for the discovery of domains of tissue microanatomy and architecture using graphs of cellular interactions. Intensity values and cellular segmentation masks are used to derive an expression matrix containing the intensity of each marker in each cell, and a graph of physical cellular interaction based on proximity which can be represented as a binary adjacency matrix. Message passing (described in b) combines the expression and adjacency matrices into a new matrix of spatially aggregated expression values which serves as the input for clustering methods. The resulting clusters represent domains of tissue microanatomy underlying the tissue architecture. The procedure can be performed jointly across several images, yielding consistent microanatomical domains across images. **b**) Graphical description of the message passing procedure, in which the adjacency and expression matrices are combined with the dot product. Note how in the message-passed graph, the node colors are linear combinations of the colors of the nodes with which they share edges. Each element in the feature matrix in this example depicts a vector of features.

UTAG then leverages the properties of matrix multiplication through linear algebra to combine the matrices in a procedure known as message passing (**Figure 1b**). In this, nodes of cells in physical proximity will receive a portion of the neighboring cell’s phenotypic information in a weighted manner, effectively diffusing the phenotypes into physically proximal cells determined by the adjacency matrix. The intermediate resulting spatially aggregated features therefore contain information on both cellular phenotypes and physical proximity between cells. This spatially aggregated feature matrix allows capture of microanatomical domains consisting of multiple cell types that are spatially homogeneously distributed. For example, arteries consist of a layer of endothelial cells surrounded by smooth muscle cells. Through message passing, endothelial cells become more like adjacent muscle cells and vice versa, effectively grouping cells with different phenotypic features based on their spatial distribution. Finally, this matrix is clustered using standard modern algorithms such as Leiden^22^ or Phenotyping by Accelerated Refined Community-Partitioning (PARC)^23^ clustering to derive domains of tissue structure in images (**Figure 1a,** orange area). In this process, the number of captured domains is determined by a customizable resolution hyperparameter which controls the coarseness in both Leiden and PARC clustering (**Extended Data Figure 1b**). Biological interpretation of the discovered domains remains however dependent on the user by contextualization in terms of their cell type composition, frequency of cellular interactions, or association with target variables such as clinically relevant outcomes. We provide a software package with the implementation, documentation, and tutorials on the application of UTAG to various datasets (https://github.com/ElementoLab/utag).

### UTAG uncovers microanatomy and principles of organ organization in the healthy lung

We first tested UTAG on healthy lung tissue images. The human lung is a highly compartmentalized tissue where the organ physiology dictates an intricate interplay between cells and matrix to create functional structures such as the airway lumen, alveolar airspace, and blood vessels. We applied UTAG to a dataset of 26 highly-multiplexed IMC lung images from three donor lung specimens, consisting of 28 markers, with a particular focus on airways extending from proximal bronchi and succeeding divisions to terminal and respiratory bronchioles (**Figure 2a**, first column, **Extended Data Table 1**). Importantly, in this dataset, each image has been manually annotated with organ-specific microanatomical domains such as airways, connective tissue, submucosal glands, vessels, alveolar space (**Figure 2a**, fourth column). The annotated structures effectively serve as a reference for microanatomical annotation of the lung. In addition, the cells in these data had been phenotyped into seven broad clusters of cell identity (**Figure 2a**, second column), which can be helpful to interpret the composition of the domains, albeit not used by UTAG.

**Figure 2:**
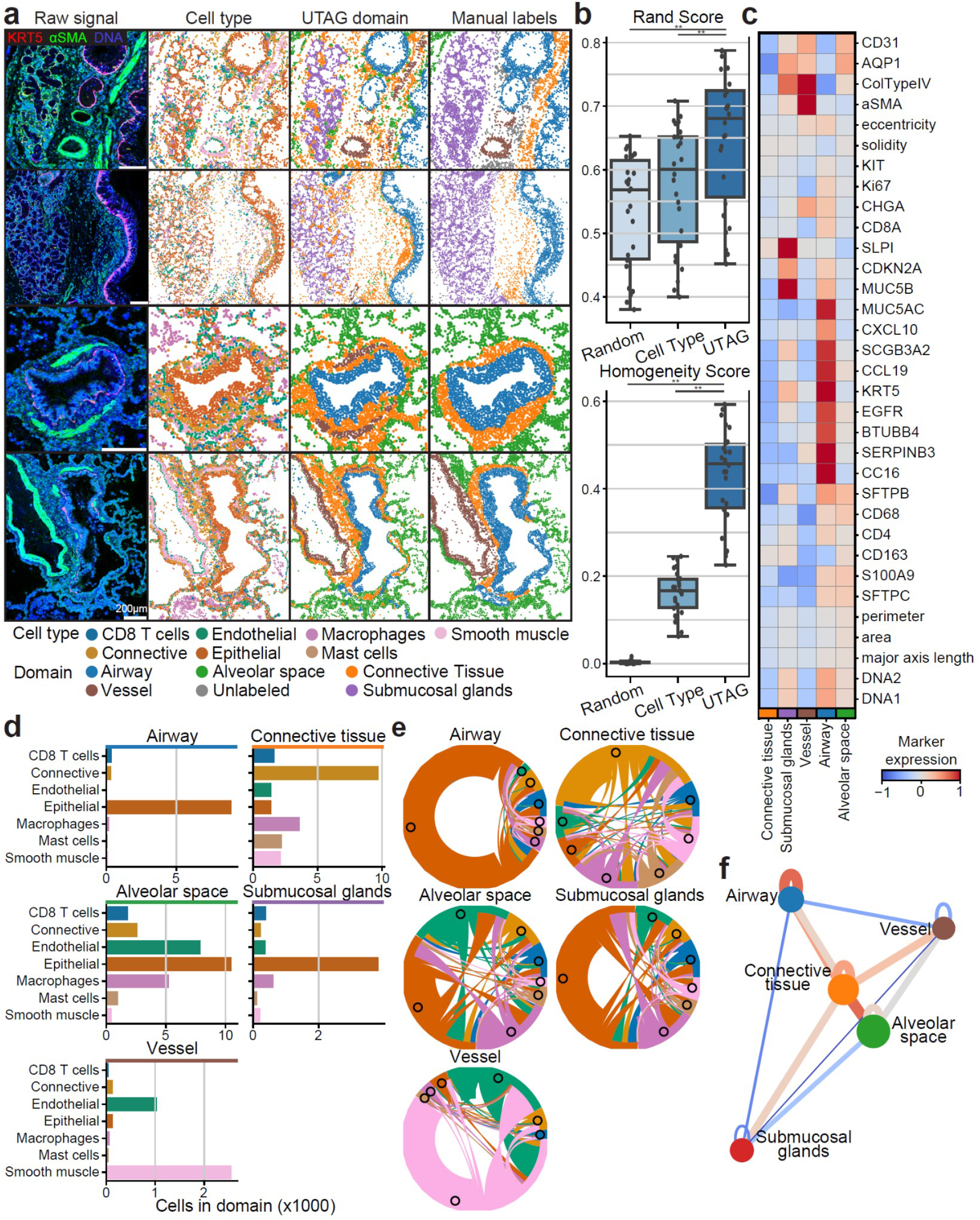
Discovery of microanatomical domains and principles of tissue architecture in human lung. **a**) microanatomical domains detected in IMC images of healthy human lung tissue. The first column illustrates the intensity of three selected channels in four representative images, the second column the cell identity of the cells in those images, the third column displays the microanatomical domains discovered with UTAG, and the fourth column microanatomical domains manually annotated by experts. Scale bars represent 200 μm. **b**) Benchmark of the UTAG domains against expert annotation, in comparison with randomized domain labels per cell, and cell type identities. Each point represents one image and for both metrics values closer to 1 are optimal.** p < 0.01, two-sided Mann-Whitney U-test. **c**) Mean channel intensity for all channels aggregated by the discovered microanatomical domains. **d**) Cellular composition of microanatomical domains. **e**) Composition of microanatomical domains in terms of intercellular interactions derived from physical proximity. **f**) Model of physical proximity between microanatomical domains in the lung. The nodes of the graph represent the microanatomical domains, and the color of the edges between them show the strength of their physical interactions. The node position is determined based on the edge weight using the Spring force-directed algorithm.

We applied UTAG to the IMC data, by providing the position of the cells in the image, and the intensity of each marker in each cell to the algorithm. We then labeled the resulting clusters with identities into five groups depending on the intensity of markers and cellular composition (**Extended Data Figure 1c**). The resulting microanatomical domains detected by UTAG largely recapitulated the microanatomy of manually labeled domains (**Figure 2a**, third column). To assess the performance of our method, we compared the discovered microanatomical domains with the annotations from expert labels using two different metrics based on cell domain properties (**Figure 2b**). As comparison, we calculated the same metrics based on randomly shuffled domain labels and cell type identities. UTAG significantly outperformed both in terms of label agreement (Rand score) and purity (Homogeneity score) (**Figure 2b**, *p* < 8.9×10^-9^, two-tailed Mann-Whiteney U-test), which shows that an unsupervised approach can discover accurate microanatomical domains in multiplexed imaging data.

The domains discovered by UTAG were enriched in protein expression specific to each domain, as evidenced by KRT5, CC16, SCGB3A2, MUC5B, and MUC5AC expression in airways, or CD31, alpha smooth muscle actin (aSMA) and type IV collagen in vessels (**Figure 2c**). Furthermore, we found the cell type composition to reflect the captured domains. Airways and submucosal glands consisted predominantly of epithelial cell types while being spatially distinct. Connective tissues were generally composed of sparse matrices of cells that had low expressions of all markers (**Figure 2c**), but sometimes included supportive muscles and infiltrating immune cells (**Figure 2d**). Other identified domains were well balanced in terms of cell type composition. The alveolar space included a well-balanced proportion of epithelial and endothelial cells required for gas exchange (**Figure 2d**). This reveals that UTAG, without specific training, is capable of effectively capturing both simple domains with a dominant cell type, and more complex domains composed of multiple cell types. Beyond cell type composition, we identified distinct differences in the frequency of physical interactions between cells of different cell types across microanatomical domains (**Figure 2e**). In airways, we observed a tight connection between epithelial cells, and reciprocal proximity between epithelial and connective tissue. The connective tissue, as a transition tissue between airways and other functional domains in the lung showed high diversity and balance in cellular interactions. The alveolar space domain has strong reciprocal interactions between epithelial and endothelial cells which is a hallmark of alveolar type 1 cells closely connected to capillary endothelium. Taken together, the observed cell type abundance (**Figure 2d**) and interaction relationships (**Figure 2e**) within the microanatomical domains of the lung provide a signature of the architecture of the healthy human lung.

While the composition of an organ in microanatomical domains is an important part of its architecture, it is also important to understand the wider-scale architecture of an organ in relation to its physiology. To demonstrate how UTAG can be useful in uncovering organ-specific high-level architecture, we quantified physical interactions between microanatomical domains in IMC images, and related domains based on the frequency of interactions (**Figure 2f**). The resulting network, made by associating the strength of microanatomical domain interaction with attraction between nodes, summarizes the architecture of the lung - with a main anatomical axis of high order tissue assembly from airway, connective tissue to alveolar space (**Figure 2f**). Furthermore, we also found that both vessels and submucosal glands, while interacting with similar domains, are diametrically opposed to the main axis (**Figure 2f**), which may suggest that segregation of vascular and secretory domains of the lung is a hallmark of healthy lung architecture. Overall, the microanatomical domains detected by UTAG in the lung, along with the inferred high-level structure of the organ illustrate the accuracy and utility of UTAG in understanding tissue architecture at various scales with a completely unsupervised approach.

### UTAG captures changes in microanatomical domain composition and structure in diseased lung tissue

Having established the performance and usefulness of UTAG in multiplexed imaging of healthy tissue, we sought to determine whether UTAG is able to discover microanatomical domains in disease as well. To that end, we ran UTAG on a dataset of 239 IMC images with 37 markers from 27 deceased patients due to lung infection^24^ (**Figure 3a**). Despite using a different set of markers from the healthy lung dataset (**Figure 2a**), we were able to discover six largely similar microanatomical domains which were present in images of various disease groups: one domain representative of epithelial cells (predominantly airways), one domain of fibroblast-rich connective tissue, one domain for alveolar regions, one for vessels, one with clusters of various immune cells, and a rare one of clustered neutrophils exclusively. Their relative abundance however, reflects the changes in the morphology and cellular composition of the tissue after infection^24^, with for example an increased proportion of the epithelial domain following Influenza and in late COVID-19, and an increase in the fraction of connective tissue in late COVID-19 indicative of fibrosis (**Figure 3b**). Since topological domains aggregate spatially proximal cells of various cell types that contribute to tissue function, we hypothesized that the abundance of topological domains across images more easily explains the variance in the dataset than the abundance of cell types on their own. Indeed, in a Principal Component Analysis reduction of the data, we found that not only the fraction of variance in the first component was higher with topological domains, but they also more easily reconstructed the linear progression of healthy tissue in comparison with cell type identities alone (**Figure 3c**).

**Figure 3:**
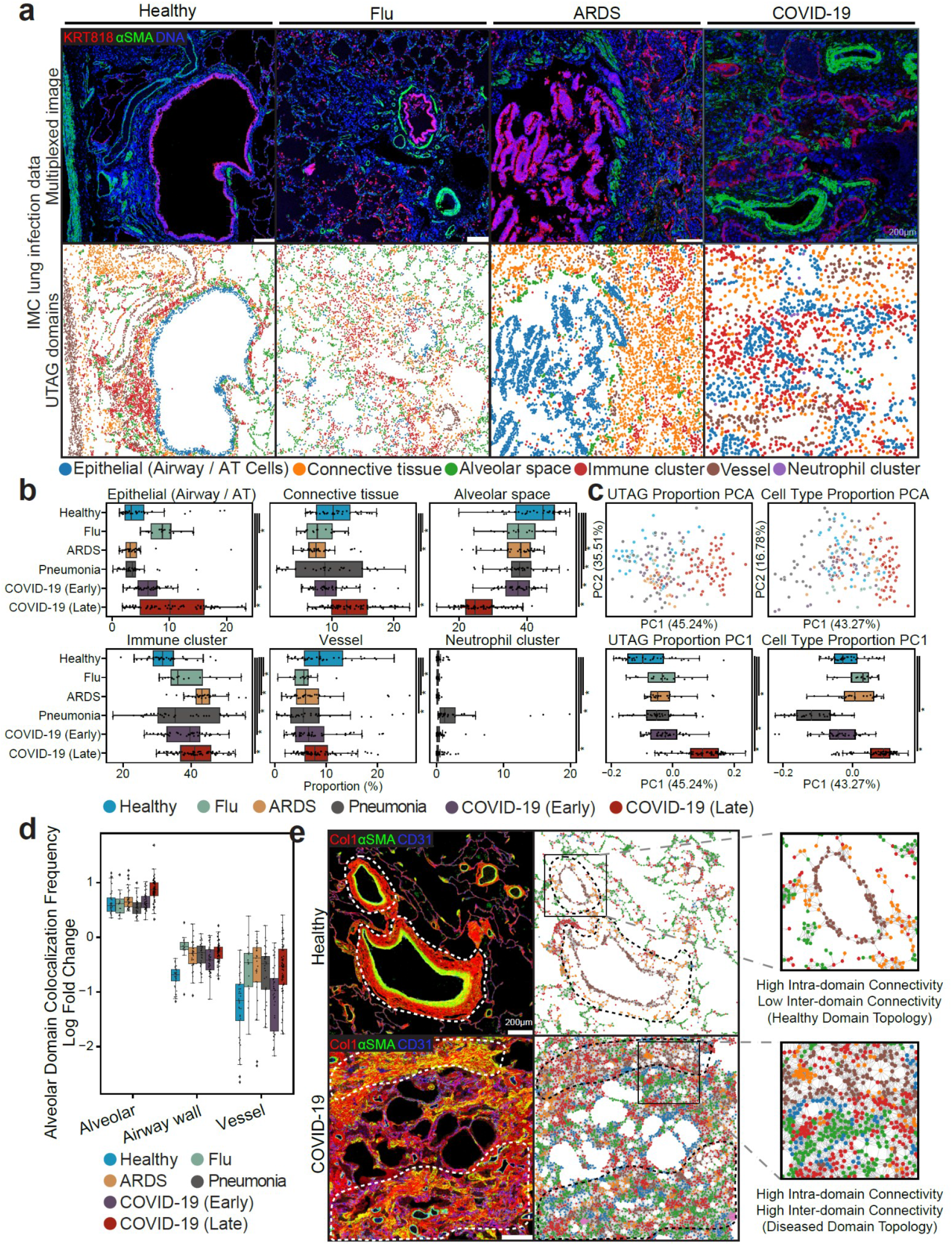
Microanatomical domains discovered by UTAG across data types and disease states. **a**) Discovery of microanatomical domains in IMC images of lung from patients of various pathologies. The top row illustrates the intensity of three selected channels, while the bottom row displays the UTAG domains. **b**) Univariate analysis of microanatomical domain composition across lung infection disease. Microanatomical domain composition was percent normalized per slide. Mann Whitney U test was used for statistical comparison between healthy and disease types. **c**) Principal Component Analysis for joint analysis of domain (left) or cell type (right) composition per image. The top two plots visualize the position of images in the first two principal components. The bottom two plots show the distribution of the first principal component aggregated by disease group. **d**) Log odds of domain colocalization frequencies across diseases in alveolar domains. Log odds indicates observed frequency over expected as estimated empirically by random permutation. Positive values indicate high intra-domain (alveolar-alveolar) colocalization compared to random mixtures, and negative indicates low inter-domain colocalization. **e**) IMC images of healthy and COVID-19 infected lung tissue. The healthy shows highly compartmentalized domains, particularly in the vasculature, while diseased lung shows loss of compartmentalization. For b) and c), asterisks represent p-values less than 0.05 after Benjamini-Hochberg adjustment.

Since differences in cell type composition during lung infection have been reported^24^, we sought to investigate whether there are differences in the high-level composition of tissue, as quantified by the spatial proximity in topological domains across images (**Extended Data 3**). The most prominent differences in topological domain colocalization between disease states was observed between the alveolar space and vessel domains (**Figure 3d**). In Influenza, ARDS and late COVID-19, vessel domains interact with alveolar domains more tightly than in Healthy lung or early COVID-19. In healthy lung sections, vessels often have high intradomain connectivity and are isolated from other domains, while in late COVID-19 we observed high connectivity of vessels with other domains, particularly the alveolar space (**Figure 3e**). This likely reflects the previously described increase in vasculature due to pathology induced angiogenesis ^25,26^. The characterization of microanatomy across various disease states in the lung along with the discovery of changes in the connectivity of tissue domains, demonstrate the versatility of unsupervised approaches such as UTAG to detect and quantify microanatomical structure in human tissue.

### UTAG uncovers tumor microenvironment domains associated with patient survival

We have so far employed UTAG in the lung because we have annotated images allowing us to assess whether the discovered microanatomy aligns with current knowledge in the field. Given that UTAG is an unsupervised method, it is not guaranteed that its use across organs, data types, and disease states will always discover microanatomical domains with physiological relevance or of pathologic interest.

To address this question, we first applied UTAG to a dataset of 19 cyclic immunofluorescence (CyCIF) images with 26 markers from 3 lung cancer patients^27^ (**Extended Data 4**). We observed that the obtained domains largely reflected tumor or stromal microenvironments reflecting a complete departure from the tissue architecture seen in normal lung. This is likely due to proliferation of neoplastic cells which is independent from the normal physiological function of the lung. In this setting, UTAG may be of use in cancer by detecting the interface between tumor and stromal, facilitating the investigation of cellular composition and interactions at this interface, without the need for manual annotation of images by an expert.

Second, to assess whether UTAG is capable of discovering microanatomy in other organs, we apply it to a dataset of 58 IMC images with 28 markers from 7 patients of upper tract urothelial carcinoma (UTUC)^28^ (**Figure 4a**). In line with our observations in lung cancer (**Figure 3c**), the five discovered domains largely reflected the division between tumor and stroma microenvironments. However, we did notice a gradient between the two, with domains with considerable immune infiltration for both tumor and stroma, and a domain present mostly at the interface between tumor and stroma (**Figure 4a**). Of note, in this dataset, 16 images had been manually annotated with boundaries of tumor and stroma, which allowed us to assess the performance of UTAG in the delineation of these boundaries (**Figure 4b**). We found that UTAG domains largely recapitulate these annotations and significantly outperformed randomly shuffled labels in both agreement and purity (**Figure 4b**, *p* < 3×10^-7^, two-tailed Mann-Whitney U-test), and cell type annotations in terms of agreement with manual labels (**Figure 4b**, *p* = 1.4×10^-3^).

**Figure 4:**
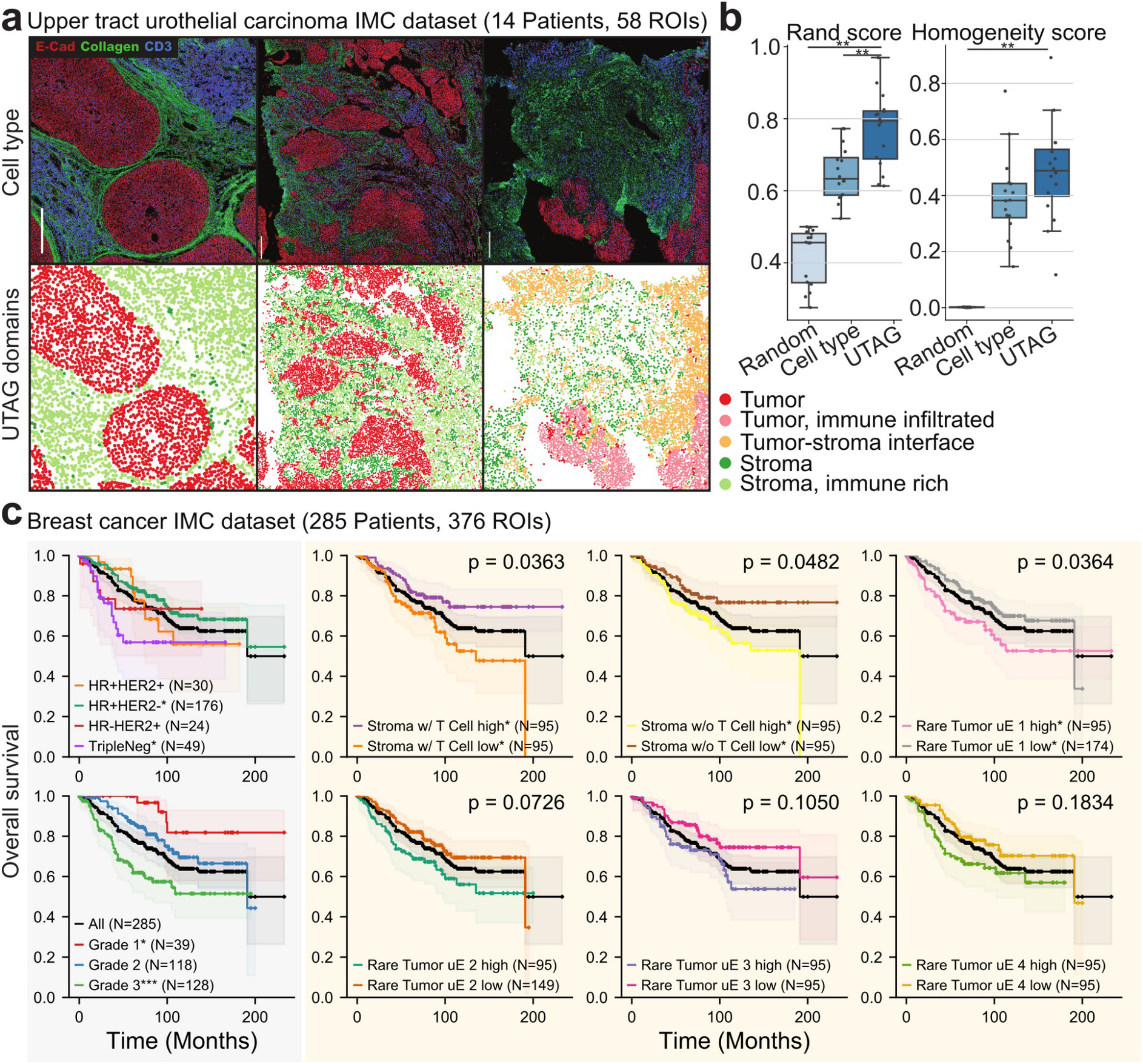
Discovery of microanatomical domains associated with cancer survival. **a**) Discovery of tumor and stromal domains in IMC images of UTUC. The top row illustrates the intensity of three selected channels, while the bottom row displays the UTAG domains.Scale bars represent 100μm. **b**) Benchmark of the UTAG domains against manual annotation of tumor and stromal domains. For comparison, we include randomized domain labels per cell, and cell type identities. Each point represents one image and for both metrics values closer to 1 are optimal. **c**) Discovery of microanatomical domains in IMC images of breast cancer and their association with overall patient survival. The first column, included for comparison, are patient survival curves depending on tumor subtype and grade classification. The remaining columns are UTAG domains for which patients with presence of the domain above the median show greatest association with survival. p-values were calculated with a log-rank test and adjusted with the Benjamini-Hochberg FDR method.

Third, we sought to assess whether domains discovered by UTAG in large datasets of multiplexed images can be associated with relevant clinical outcomes of patients. We used a dataset of 376 IMC images with 32 markers from 285 breast cancer patients^15^, with associated clinical data on tumor subtypes, staging, and overall survival (**Figure 4c**, first column). We ran UTAG on the dataset and discovered 30 types of microanatomical domains (**Extended Data 5a**) including tumor and stromal regions. We then quantified the presence of each domain in each patient, and divided patients in tertiles based on the abundance of that domain across all samples. By comparing patients in the top vs bottom tertiles, we found 3 domains where the abundance was significantly associated with overall survival (**Figure 4c** and **Extended Data 5b**, p < 0.05 log-rank test with Benjamini-Hochberg FDR correction).

Two of these associations were in largely complementary domains, related with the abundance of T-cells in stromal components, where low abundance of T-cells is associated with lower overall survival (**Figure 4c**). This could be due to a potential immune action against the cancer cells by T-cells.

In summary, our analysis of tumor microenvironment domains in large cohorts of cancer patients revealed the accuracy of UTAG in detecting microenvironments reflecting tumor/stromal boundaries in agreement with manual annotations, and shows that unsupervised discovery of tissue domains can have prognostic value in disease for example in patient stratification.

## Discussion

In this study we develop UTAG, a novel method to perform discovery and quantification of microanatomical tissue structures in biological images which uses no prior knowledge. Our method leverages the combination of phenotypic and proximity information of cells to discover topology of tissues in various organs, and various types of multiplexed imaging data. Given the lack of formal definition of microanatomical domains and healthy tissue datasets with such annotation which can be used as ground truth, benchmarking of our method relied on two datasets of lung microanatomy and tumor/stroma divisions in cancer. While we observed that UTAG performed better than random or simple cell type identities in the discovery of tissue microanatomy, the ground truth set of manual annotations is inherently subjective to the observer, and often incomplete by focusing on a subset of specific predefined structures. In fact, it is conceivable that a fully unsupervised method such as UTAG is able to capture gradients of mixtures between known domains or even novel or poorly defined structure in tissue that is underappreciated.

On top of its ability to detect tissue architecture, UTAG can serve as a method to quantify biologically relevant processes such as angiogenesis in native tissue conformation. In this paper, we presented ways to numerically quantify the loss of compartmentalization of vessels in alveolar space of COVID-19 infected lung (**Figure 3d**). In similar fashion, UTAG can be used to quantify the extent of various biological processes such as angiogenesis in individual samples - just as existing computational methods based on genomics and transcriptomics can, but with the advantage that the manifestation of biological processes are directly observable in the original physical context of the tissue.

While we believe our method provides a significant step toward the systematic discovery of tissue structure, one crucial aspect for its successful application is the interpretation of the discovered topological domains in terms of their identity and biological relevance. We demonstrated how on cases such as healthy tissue with well defined structure related with organ-specific physiology, interpretation of domain identity based on cell type composition and interactions can be achieved (**Figure 2**), while in tissues without strong structural patterning, or with undefined function such as cancer, interpretation of discovered domains can rely on the association with clinically relevant outcomes (**Figure 4**). UTAG provides flexibility to the user to discover structures present in biological images, but we believe that its potential is maximized by the involvement of experts in the field such as pathologists, in the discovery process and interpretation of results.

Beyond the conceptual limitation in the biological interpretation of UTAG results, a few technical issues must also be taken into account. UTAG relies on user-supplied cell segmentation to determine positional information from the cells and consequently infer physical interactions. Recent advances in cellular segmentation algorithms^29–32^ have greatly advanced the quality of segmentation masks for various types of images, but downstream results can only be as good as the segmentation. Furthermore, we greatly simplify the geometric complexity of 2D tissue slices by assuming centroids capture most of the positional information of cells, which for eccentric cell types such as neurons, endothelial, and various types of eccentric immune cells, may not be the case.

The inference of cellular contacts and the scale at which microenvironmental signals diffuse across the local cellular context are fields of current study^33–36^ and of importance for the detection of tissue microanatomy. UTAG requires a user-provided parameter to discretize cellular contacts. In our experience, we found that changes in this parameter were most needed depending on the resolution of the images, since optical imaging typically has more resolution than for example laser-based tissue ablation in IMC. Nonetheless, this is something we purposefully designed to be tuned by the user in order for UTAG to be adaptable to the requirements of various image types and cellular contexts without assumptions on the underlying structure of the tissue, such as has been done for example relying on the consistent shape of germinal centers across organs^37^.

UTAG opens new possibilities in our ability to understand tissue architecture by detecting microanatomical domains, but also by quantifying how they interact at a higher level to a point that we could infer the broad rules of human lung architecture. We envision that in the future, UTAG could be applied to traditional hematoxylin and eosin stained histopathological images if an appropriate feature matrix of stain intensity and cell morphology can be extracted. That would open the possibility for the detection of microanatomical structures in large biobanks and association of these with clinical features at scale. Likewise, systematic application of UTAG in image data from various organs will undoubtedly accelerate projects such as spatial cell atlases^38–40^, by providing microanatomical context to the cells, and enabling ground-up discovery of tissue architecture beyond cell type composition of tissues. Another exciting future application is the discovery of microanatomy in volumetric images of tissue^13,41–43^, since there is no conceptual limitation to use UTAG in 3 dimensions. This would enable robust morphometry of tissue structures since a current challenge in two-dimensional analysis of tissue is the detection of structure independent of the cutting angle. Robust, automated assessment of tissue microanatomy could enable the definition of tissue integrity ranges in healthy human tissue across ages, detection of early pre-cancer lesions and cancer invasion, and the study of age-associated diseases characterized by cellular degeneration, fibrosis, and loss of tissue integrity such as chronic obstructive or idiopathic pulmonary disease.

## Methods

### UTAG Algorithm

The two inputs to the UTAG algorithm were the cell feature matrix and the location matrix. The cell feature matrix is designed to be as generalizable as possible to incorporate multiple imaging modalities and can contain any features ranging from generic cell properties such as cell area, perimeter, and morphology to modality specific attributes such as intensity of hematoxylin and eosin from H&E staining to marker expression quantification such as CD4, KRT8, or PD1 levels in IMC. From the location matrix, we build a graph using squidpy^44^ (version 1.1.0) where each node is a unique cell and edge is whether two cells are within a threshold Euclidean distance. We then perform message passing, an inner product between the adjacency matrix of the graph and the feature matrix, so that each cell within the graph inherits features from its immediate neighbors. When aggregating spatial components with the feature matrix, we optionally add a normalization step to allow users to take the L1 norm of the adjacency matrix so that the users can decide whether to mean-aggregate or sum-aggregate. We denote the resulting matrix spatially aggregated feature matrix that encodes information of both single cell features and cell locations. The cells in the spatially aggregated feature matrix are clustered into groups using the Leiden^22^ (version 0.8.7) and PARC^23^ (version 0.31) algorithm at multiple resolutions. Each cluster can then be annotated into microanatomical domains based on enrichment profiles or by inspecting user provided cell type identities.

### Running UTAG on IMC data

To quantify cellular phenotypes, we used the cell masks, and aggregated all pixels of a cell with the mean intensity for each IMC channel. We combined the per cells expression vector from all cells in all images into a single matrix. We then performed log transformation, Z-score normalization with truncated at positive and negative 3 standard deviations, followed by Combat^45^ (version 0.3.0) batch correction to phase out sample-specific biases. This was subsequently followed by a final Z-score normalization truncated at 3 standard deviations.

For the healthy lung dataset, UTAG was run with a *max_dist* of 15, which are in physical dimensions 20 microns (**Extended Data 1b**). For all other IMC datasets (lung infection, UTUC, and BRCA), we ran UTAG with *max_dist* of 20. Each dataset was clustered at resolutions of 0.05, 0.1, 0.3, and 0.5. The principle of selecting the optimal resolution was based on how diverse each dataset was, or in other words how many patients and diseases each dataset contained. Higher resolutions, resulting in more clusters, were preferred in diverse datasets while more homogenous datasets required only a few clusters. For the normal lung dataset, we used Leiden clustering at 0.3 resolution and annotated the resulting 11 clusters into 5 microanatomical domains (**Extended Data 1c and 1d**). For the infected lung, UTUC, and BRCA dataset we used PARC clustering with resolution 0.3, 1.0 and 0.1 respectively which resulted in 20, 34, and 31 clusters.

### Benchmarking against manual expert annotation

To show that the gain of information using the UTAG algorithm is statistically significant, we compare cell types and UTAG labels against manual expert annotations. To objectively assess the performance of UTAG, we used Rand score and homogeneity score as an evaluation metric for the unsupervised segmentation task. Rand score, also known as Rand index, is a similarity measurement that is calculated by the ratio of agreeing pairs over all pairs between the predicted and true labels. The homogeneity score^46^ assesses how uniquely predicted labels associate with true labels (a measure of cluster purity). Ranges of both metrics are from 0.0 to 1.0 inclusive, with higher scores indicating better performance. To lay out a baseline for how the metrics works, we show how random labels perform against the expert annotation. To test for differences in performance, we perform a two-tailed Mann-Whitney test between random labels scores, cell-type scores, and UTAG scores.

### Quantification of cellular interactions and microanatomical domain interactions

As UTAG achieves microanatomical domain annotation based on graphs leveraging spatial proximity, we can take advantage of the spatial neighborhood information for downstream analysis. Under the graph formalism, we can quantify cellular and domain interactions from edge counts connecting distinct nodes, identified by cell type and domain properties. Graphs were constructed with a threshold distance of 40 pixels for healthy lung IMC samples to allow a more lenient interaction threshold compared to the UTAG default. For cell-to-cell interactions, we quantify edges connecting a cell type to another and aggregate the connections into an adjacency matrix denoting the cell type colocalization. We present this cellular interaction matrix as a chord plot generated by holoviews python library. Microanatomical domains are similarly aggregated for each domain-to-domain interaction. These results are presented as a networkx^47^ (version 2.6.2) graph in a spring-force layout which visually demonstrates how each domain colocalizes with others. This was done on the logarithm of the counts of edge connections to ensure that the counts are on a comparable scale.

### Lung infection data univariate and Principal Component Analysis

To quantify the difference in domain composition across disease types, each IMC slide was aggregated by the number of cells in each domain. Cell counts were subsequently percent normalized to take into account the difference in cell densities. We perform a univariate domain proportion comparison for each disease group with respect to healthy samples using a two-sided Mann-Whitney U test. For a multivariate analysis, we reduce the dimensionality of domain proportion using Principal Component Analysis. We then perform a two-sided Mann-Whitney U test on the first principal component, similar to the univariate analysis, to show how all domain distributions jointly vary across disease. To show that the first principal component of domain proportions better captures the difference in diseases, we perform the same analysis with cell type proportions. All Mann-Whitney U tests were performed using pingouin^48^ (version 0.3.12) and were Bonferroni-Hochberg corrected.

### Quantification of domain colocalization frequency

Quantifying domain-to-domain colocalization by counting the number of edges may not provide the most representative measurement, because the value is largely by the original domain abundance. For example, if there is one domain that is more abundant than every other domain, then that domain generally has the highest colocalization count with all other domains. In order to compensate for the original domain distribution, we repeatedly perform domain permutation, random shuffling of domains for cells in the graph, to establish an expected colocalization frequency given the domain distribution. We add one to both the observed colocalization frequency and expected frequency, computed by the mean of 100 random permutations, to avoid division by zero. Log-fold change for domain colocalization is then computed by taking differences between two log-transformed values.

### Running UTAG on CyCIF data

40X CyCIF lung cancer samples were downloaded from DOI 10.7303/syn17865732. We used the provided cell segmentation probability maps generated with standard watershed algorithms in ImageJ or MATLAB to create cell masks using DeepCell, similar to the IMC data preprocessing. Cell fluorescence was mean aggregated just as in the IMC data. All cells across images were combined together, and the resulting matrix was log transformed, Z-scaled, batch corrected with Combat, and Z-scaled again.

Before running the UTAG algorithm, 11 DNA channels and 7 background channels were removed from the feature matrix, leaving 26 channels to remove background noise and to ensure that the algorithm was not overly influenced by replicates of a single feature. The UTAG algorithm was run with a thresholding distance of 50 pixels because the per pixel distance was more than twice as high at this magnification. We ran both Leiden and PARC clustering at multiple resolutions of 0.05, 0.1, 0.3, 0.5, and 1. We annotated stromal and tumor regions based on 0.1 resolution, as the seven created clusters were more than enough for a small dataset with 3 patients and 16 slides.

### Association of UTAG domains with survival in breast cancer

To assess whether UTAG labels have clinical implications, we run UTAG on a 285 patient breast cancer dataset with survival information. We calculate the average microanatomical domain density (number of cells/mm^2^) for each cluster created by PARC clustering at resolution 0.1. We build a domain density matrix by concatenating the domain densities for all patients. Very rare domains whose 75th percentile was zero were excluded from downstream analysis. For each domain, we categorize the 285 patients into two disjoint groups by using the two extremities thresholded by ⅓ and ⅔ quartile of domain densities (**Extended Data Figure 5**). To identify how each domain density impacts patient prognosis, we perform Kaplan Meier survival analysis using the lifelines package^49^ (*lifelines.KaplanMeierFitter*) (version 0.26.4). In order to test whether there is a statistically significant difference between the survival probability of two subgroups, we calculate p-values using a log rank test, and adjust p-values for multiple testing correction using pingouin^48^ (version 0.3.12). We also include the survival analysis results stratified by clinical hormone subtypes and tumor grades for reference.

Software used: squidpy^44^ 1.1.0, Leiden^22^ 0.8.7, PARC^23^ 0.31, ilastik^50^ 1.3.3, DeepCell^32^ 0.10.0, Combat^45^ 0.3.0, StarDist^29^, lifelines^49^ 0.26.4, scikit-image^51^, scikit-learn^52^ 0.24.2, scanpy^53^ 1.8.0, pingouin^48^ 0.3.12.

## Data availability

Healthy lung IMC data are part of a publication submitted concurrently to BioRxiv and will be made public upon publication of that work. The remaining datasets are publicly available at the following URLs:

- COVID-19 lung IMC^24^: https://doi.org/10.5281/zenodo.4110559
- Lung cancer t-CyCIF^27^: https://doi.org/10.7303/syn17865732
- Upper tract urothelial carcinoma IMC^28^: https://doi.org/10.5281/zenodo.5719187
- Breast cancer IMC^15^: https://doi.org/10.5281/zenodo.3518283

## Code availability

Source code is publicly available at the following URL: https://github.com/ElementoLab/utag

## Acknowledgements

A. F. R. is supported by a NCI T32CA203702 grant. O.E. is supported by NIH grants UL1TR002384, R01CA194547, and Leukemia and Lymphoma Society SCOR 7012-16, SCOR 7021-20 and SCOR 180078-02 grants.

## Author contributions

J.K., A.F.R., and O.E. planned the study; J.K., and A.F.R., performed analysis. S.R., and R.S. provided expertise in pulmonary biology, histology, and definition of microanatomical domains. O.E., supervised the research. J.K., A.F.R., and O.E. wrote the manuscript.

## Competing financial interests

O.E. is scientific advisor and equity holder in Freenome, Owkin, Volastra Therapeutics and OneThree Biotech. The remaining authors declare no competing financial interests.

## Extended data figures

**Extended Data Figure 1:**
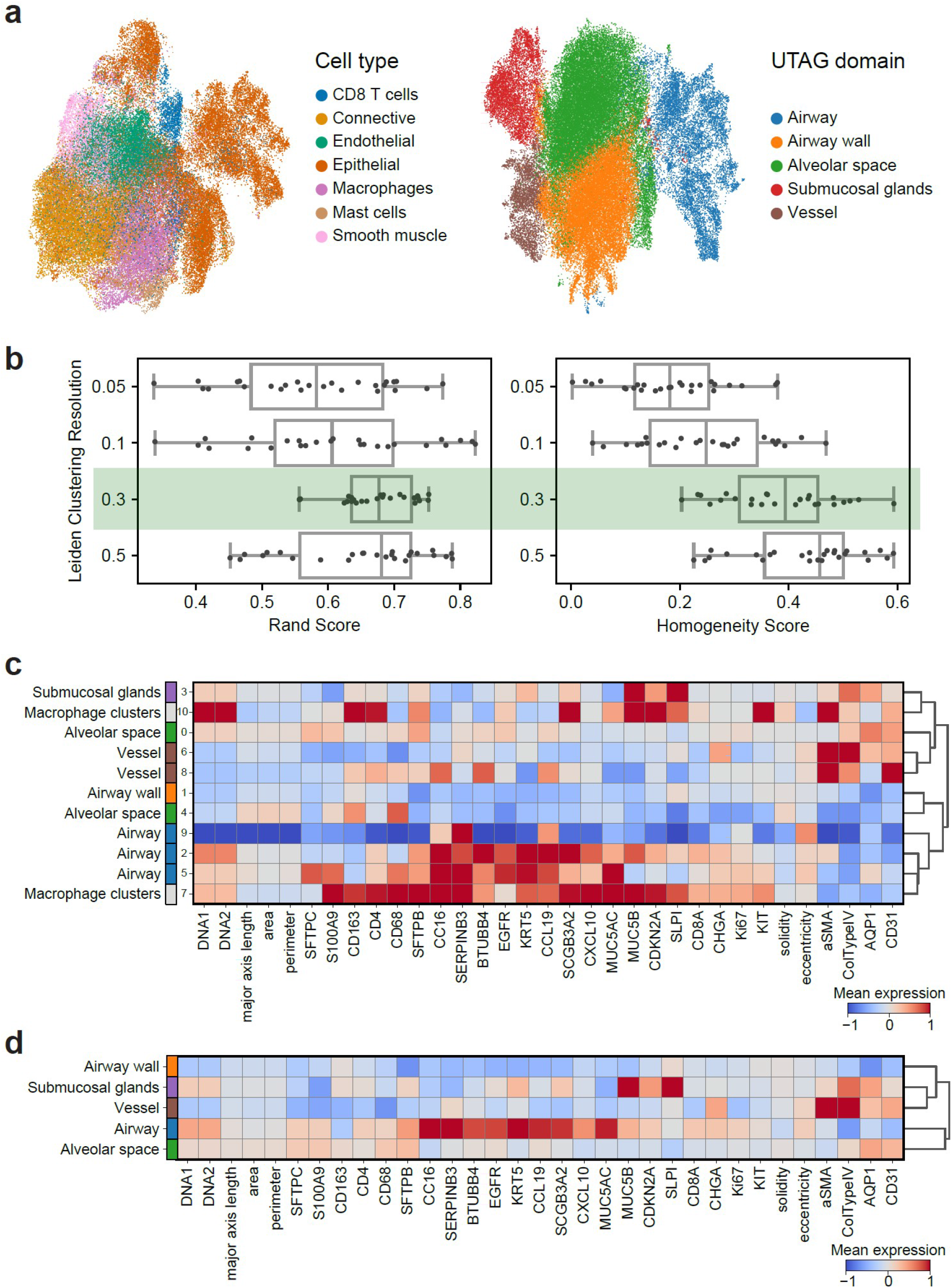
UTAG analysis of IMC images of healthy lung. **a**) UMAP representation of all cells across all images based on cellular phenotypes only (left), or cellular phenotypes and positional information combined with UTAG (right). **b**) Labeling of domains from clustering indices. Cluster indices from leiden clustering at resolution 0.1 were mapped to domains based on expression profiles. **c**) Deciding optimal resolution for healthy lung IMC data. Leiden clustering for resolution of 0.1 was selected as the ideal resolution because it had the greatest median rand score across all slides.

**Extended Data Figure 2:**
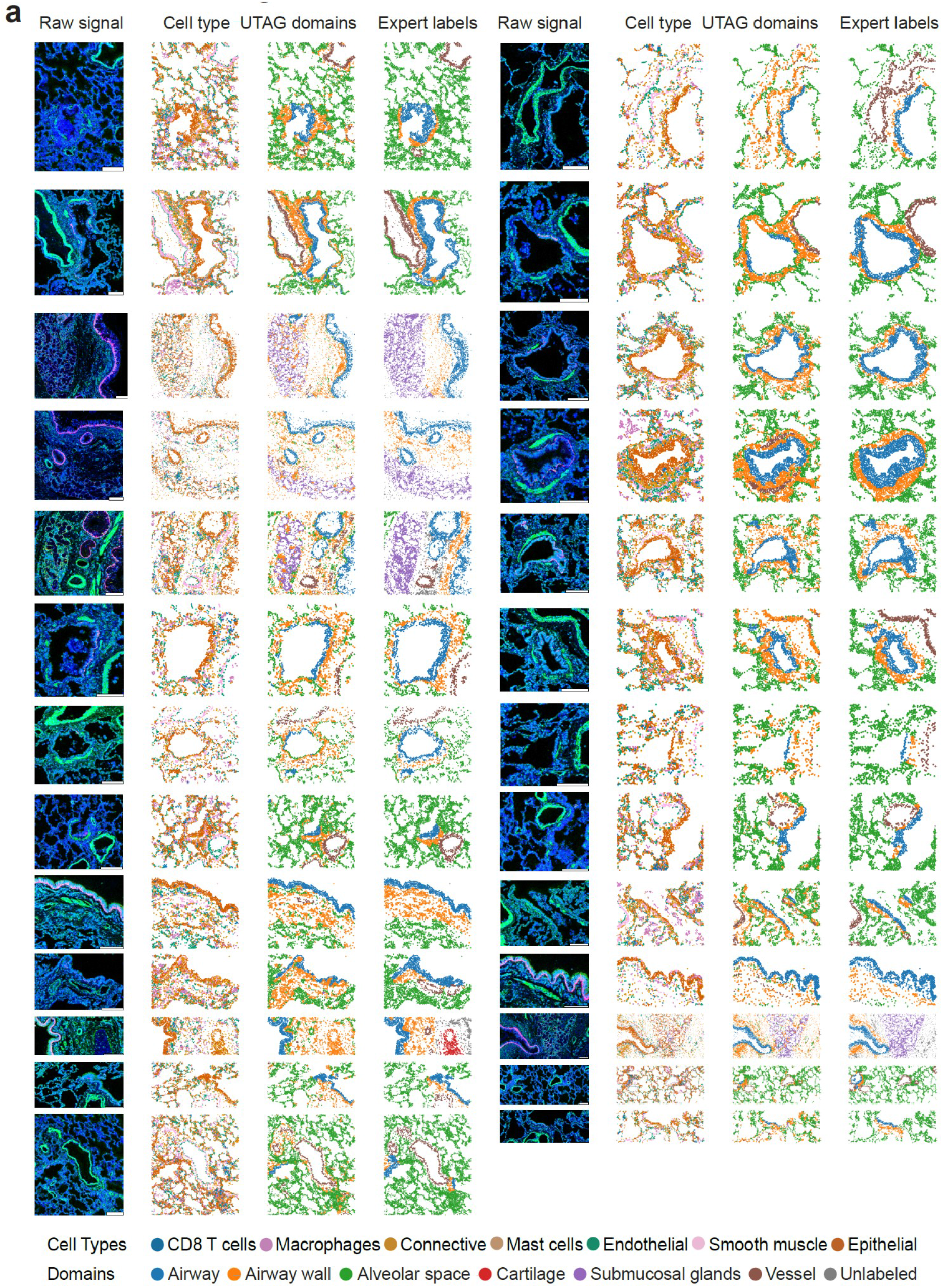
UTAG results on IMC images of healthy lung. **a**) Illustration of lung IMC images where the first column illustrates three channels (X, Y, Z), the second column cell type identities, the third column cells colored by manual annotation of microanatomical domains, and the fourth column cells colored i by UTAG domains. Each channel on the raw signal is keratin 5 for red, alpha smooth muscle for green, and DNA for blue. Scale bars represent 200 μm.

**Extended Data Figure 3:**
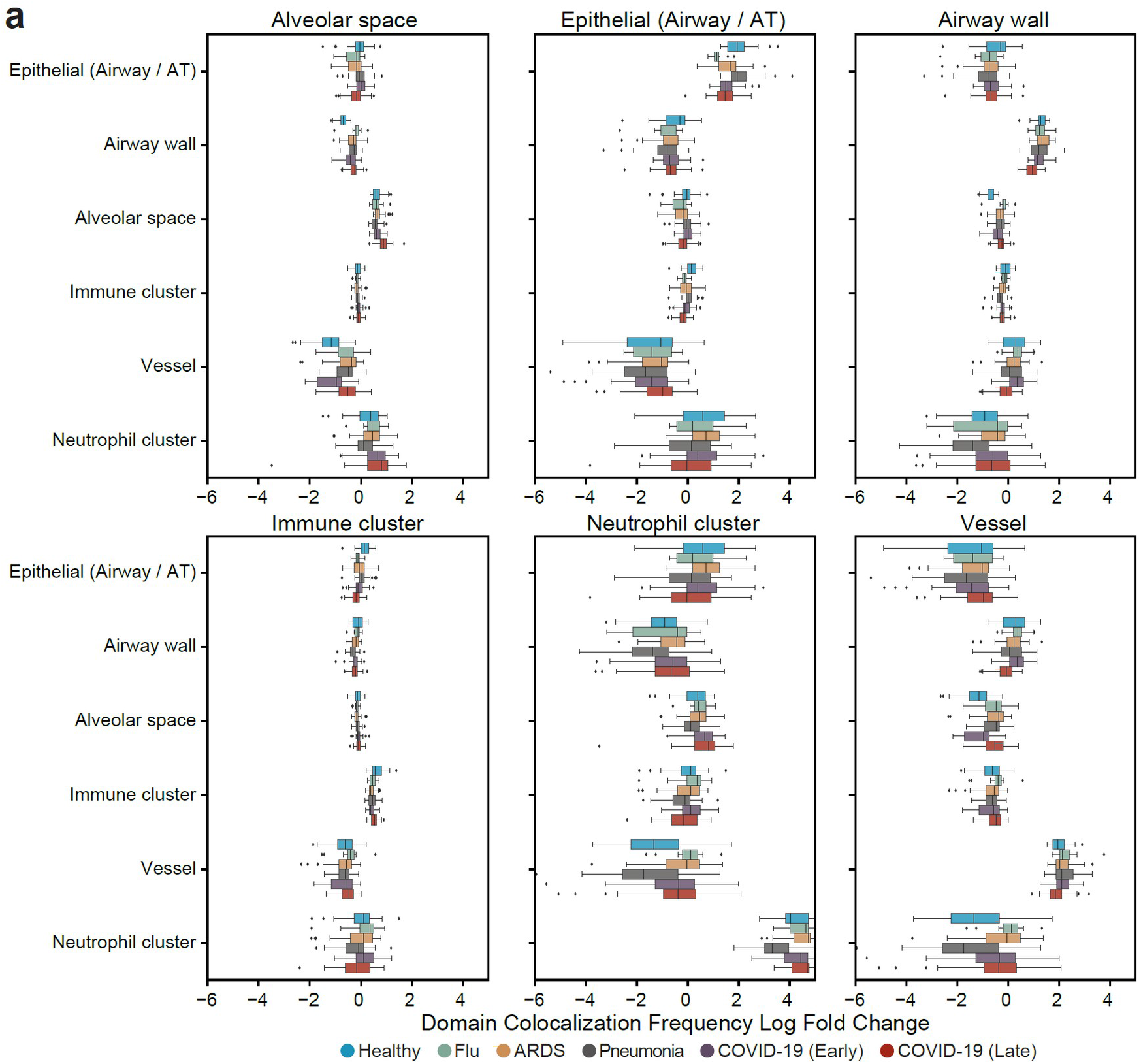
Application of UTAG to quantify domain colocalization frequency. **a**) Full comparison of domain colocalization frequency for all pairwise microanatomical domains in lung infection data grouped by disease type.

**Extended Data Figure 4:**
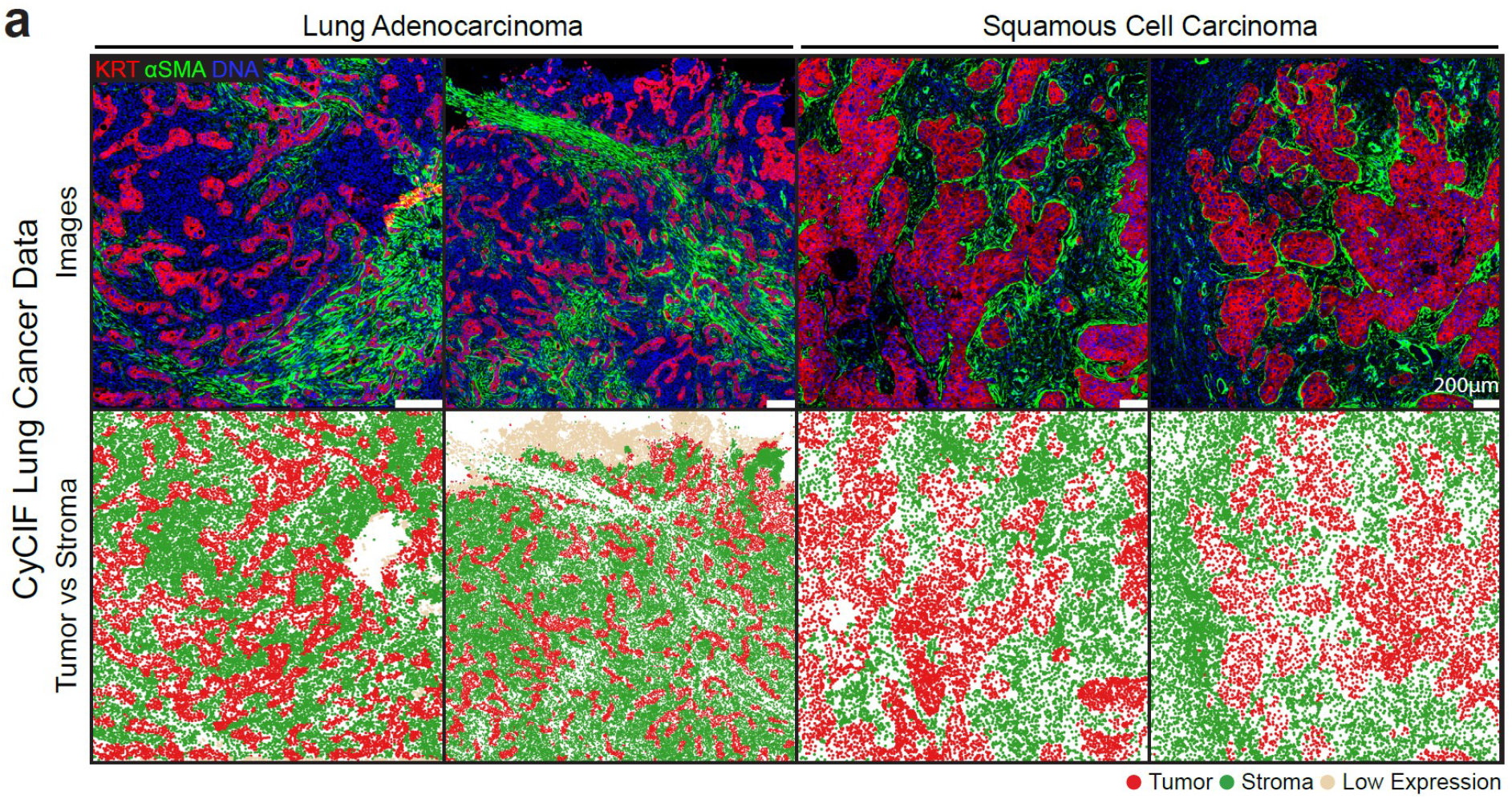
Application of UTAG to CyCIF lung cancer data. **a**) Discovery of tumor and stromal domains in CyCIF images of two types of lung cancer. The top row illustrates the intensity of three selected channels, while the bottom row displays the UTAG domains. Scale bars represent 200 μm.

**Extended Data Figure 4:**
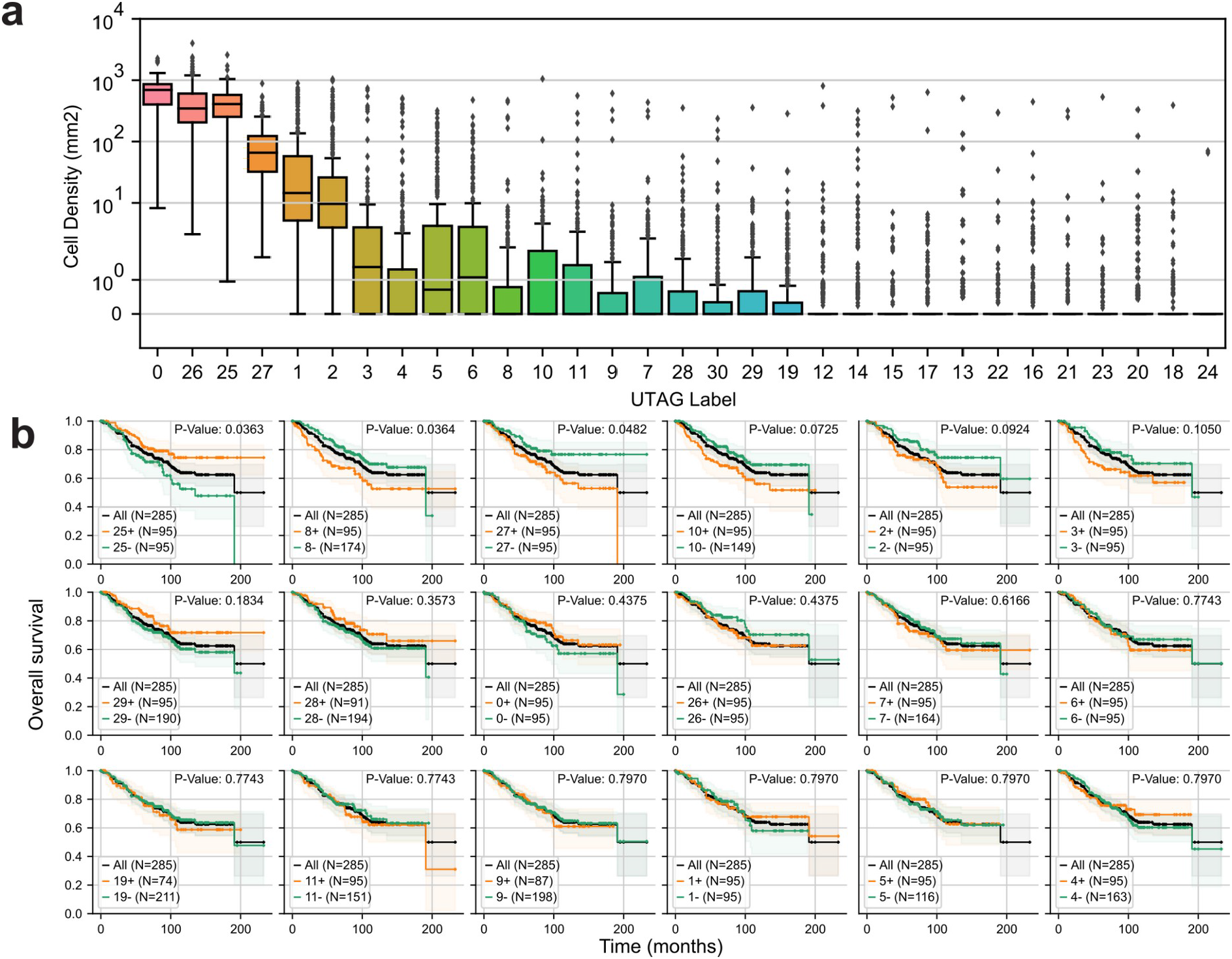
Application of UTAG to images of a cohort of breast cancer patients. **a**) Abundance of discovered UTAG clusters in each image. **b**) Survival analysis for patients based on whether they are above or below the mean abundance of each UTAG cluster. p-values were calculated with a log-rank test and adjusted with the Benjamini-Hochberg FDR method.

## Extended data tables

**Extended Data Table 1.** Samples included in analysis of healthy lung tissue with IMC.

| Sample ID | Age<sup>1</sup> | Sex<sup>2</sup> | Ancestry<sup>3</sup> | Smoking<sup>4</sup> | Lung disease | Cause of Death<sup>5</sup> | Source<sup>6</sup> | Internal ID |
| --- | --- | --- | --- | --- | --- | --- | --- | --- |
| N1 | 63 | F | C | 13 PY | No | Stroke, ICH | UNC | SR-1002 |
| N2 | 32 | M | C | 1.8 PY | No | Anoxia, cardiovascular (natural causes) | UNC | UNC-59 |
| N3 | 60 | M | A | NS | No | NA | UNC | H-9293 |

<sup>1</sup> Age (years);
<sup>2</sup> Sex: F, female; M, male;
<sup>3</sup> Ancestry: A, African-American; C, Caucasian
<sup>4</sup> Smoking: for smokers - pack-years (PY); non-smokers (NS)
<sup>5</sup> Cause of death (CoD) : ICH, intracerebral hemorrhage; NA, specific information not available
<sup>6</sup> Sample source: UNC, University of North Carolina Tissue and Cell Culture Procurement Core

## Notes

https://github.com/ElementoLab/utag

